# Impaired learning and memory ability induced by a bilaterally hippocampal injection of streptozotocin in mice: involved with the adaptive changes of synaptic plasticity

**DOI:** 10.1101/2020.08.04.235481

**Authors:** Congcong Qi, Xingxing Chen, Xinran Gao, Jingxian Xu, Sen Liu, Jinfang Ge

**Affiliations:** Department of Laboratory Animal Science, Fudan University, Shanghai, China; School of Pharmacy, Anhui Medical University, Hefei, China; Anhui Provincial laboratory of inflammatory and immunity disease, Anhui Institute of Innovative Drugs, Hefei, China; The Key Laboratory of Anti-inflammatory and Immune Medicine, Ministry of Education, Anhui Medical University, Hefei, China

**Keywords:** Alzheimer Disease, Streptozotocin, BDNF, Nesfatin-1, NMDAR

## Abstract

**Background:** Alzheimer’s disease (AD) is a neurodegenerative disease characterized by progressive cognitive decline, psychiatric symptoms and behavioral disorders, resulting in disability and loss of self-sufficiency.

**Objective:** To establish an AD mice model, investigate the behavioral performance, and explore the potential mechanism.

**Methods:** Streptozotocin (STZ, 3 mg/kg) was microinjected bilaterally into the dorsal hippocampus of C57BL /6 mice to establish the AD model. Behavioral changes (anhedonia and despair, balance and motor coordination, locomotion, and learning and memory) were examined and the serum concentrations of insulin and nesfatin-1 were measured by ELISA. The activation of hippocampal microglia was assessed by immunohistochemistry and the protein expression of several molecular associated with the regulation of synaptic plasticity in the hippocampus and the prefrontal cortex (PFC) was detected via western blotting.

**Results:** The STZ model mice showed a slower bodyweight gain and higher serum concentration of insulin and nesfatin-1. Although there was no significant difference between groups with regard to the ability of balance and motor coordination, the model mice presented a decline of spontaneous movement and exploratory behavior, together with an impairment of learning and memory ability. Increased activated microglia was aggregated in the hippocampal dentate gyrus of model mice. Moreover, the protein expression of NMDAR2A, NMDAR2B, SynGAP, PSD95, BDNF, and p-β-catenin/β-catenin were remarkably decreased in the hippocampus and the PFC of model mice, and the expression of p-GSK-3β (ser9)/GSK-3β were reduced in the hippocampus.

**Conclusion:** A bilateral hippocampal microinjection of STZ could successfully duplicate an AD mice model, as indicated by the impaired learning and memory and the alternated synaptic plasticity, together with the hyperactive inflammatory response in the hippocampus and the imbalanced abundance of serum insulin and nesfatin-1. Apart from these, the mechanism might be associated with the imbalanced expression of the key proteins of Wnt signaling pathway in the hippocampus and the PFC.

**Graphical abstract:** 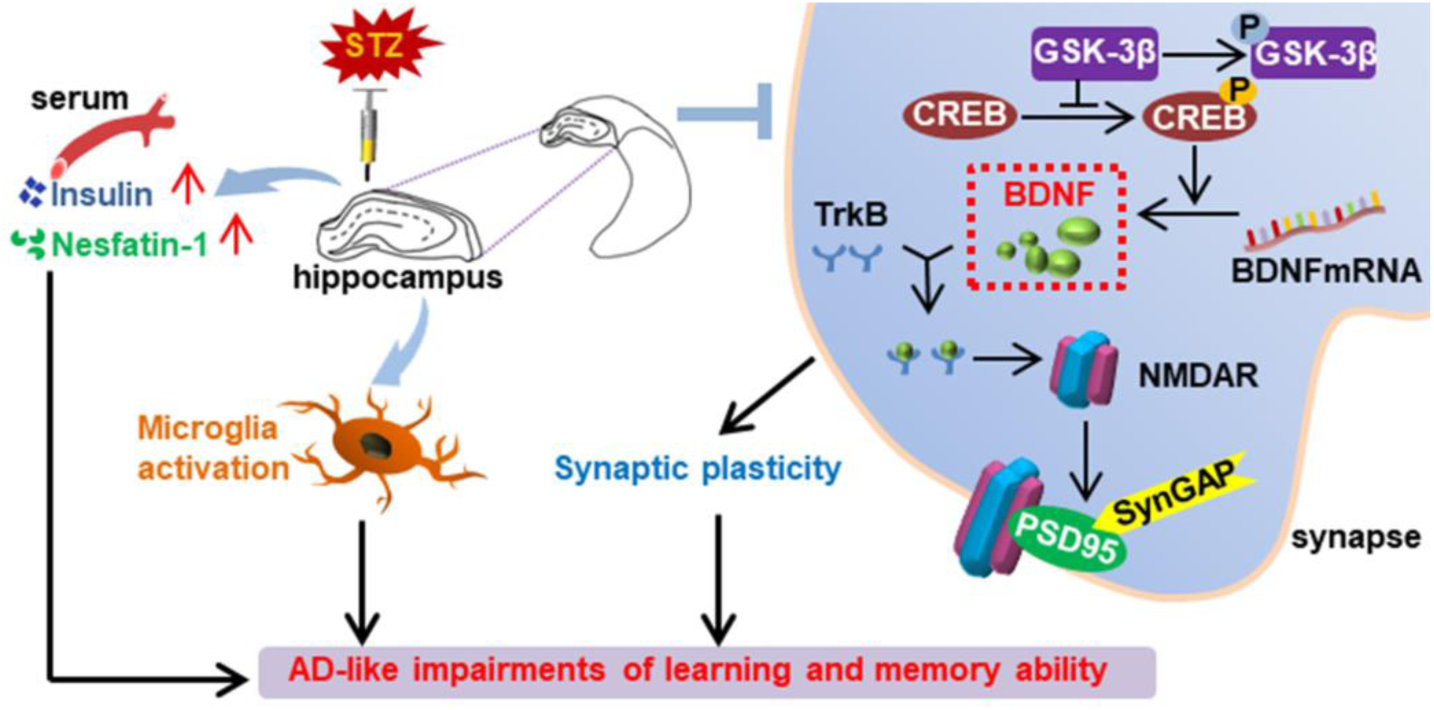

## Introduction

Alzheimer’s disease (AD) is a devastating neurodegenerative disorder characterized by a progressive deterioration of cognitive functions (Zameer et al., 2019). Targeted at increasing the activity of cholinergic neurons, cholinesterase inhibitors such as tacrine are clinically used in the treatment for AD with limited effect, but they could not slow or reverse the progress of AD. According to the classical amyloid cascade hypothesis, the deposition of amyloid beta (Aβ) is taken as the trigger of AD leading to the formation of senile plaques (SPs) and nerve fiber tangles (NFTs), resulting in the death of nerve cells and dementia eventually. However, the results of considerable clinical trials of anti-AD agents targeting at Aβ are mostly ended in failure (Hung & Fu, 2017). Thus, it is one of the most imperative scientific issues to explore the pathogenesis of AD clearly, find potential therapeutic targets, and develop effective therapeutic agents in the field of neuropsychiatric diseases and public health.

Increasing evidence show the important role of glutamate-mediated excitotoxity in the pathogenesis of AD, basing on which N-methyl-D-aspartate receptor (NMDARs/GluN) antagonist memantine has been approved by American Food and Drug Administration (FDA) in treatment for AD. Classical NMDAR is a heterotetramer composed of two GluN1 subunits and two GluN2 (A/B) combinations (Henneberger, Bard, King, Jennings, & Rusakov, 2013). Apart from regulating the transmission of neurotransmitters between synapses, GluN2B-mediated synaptic plasticity has been reported to play a seminal role in the formation of learning and memory, with different ways of GluN2A and GluN2B (Ballesteros, Buschler, Köhr, & Manahan-Vaughan, 2016; Brigman et al., 2010; C.-C. Wang et al., 2011). It has been demonstrated that the levels of phosphorylated and non-phosphorylated NMDAR subunits are reduced in the hippocampus of AD patients, and these abnormalities are associated with the abnormal presynaptic changes and cognitive deficits in AD (Sze, Bi, Kleinschmidt-DeMasters, Filley, & Martin, 2001). Although the mechanism remains unclearly, it might be partly contributed to the findings that the Aβ-derived diffusible ligands (ADDLs) could bind to NMDARs and cause abnormal calcium homeostasis, leading to oxidative stress and ultimately to the synapse loss and LTP inhibition (Pozueta, Lefort, & Shelanski, 2013).

NMDARs are reported to be involved in the pathogenesis of AD via regulating the synaptic function (Muller, Jacobi, Sakimura, Malinow, & von Engelhardt, 2018). Synaptic Ras GTPase activation protein (SynGAP) located at excitatory synapses is reported to integrate with postsynaptic density protein 95 (PSD95), Shank and Ca^2+^/CaM kinase II (CaMKII) into key signaling protein complexes. The complex is responsible for the membrane binding between synaptic terminals and the neurotransmitter receptors including NMDARs and AMPAR in the postsynaptic density (X. Chen et al., 2015). These results suggest that NMDARs perhaps take part in the etiopathology of neuropsychiatric injury in AD, with different physiological and molecular characteristics for different subunits of NMDARs.

The canonical Wnt/β-catenin signaling pathway is required for the maintainence of normal brain structure and function (Gaesser & Fyffe-Maricich, 2016), and the deficit of Wnt/β-catenin pathway has been demonstrated in the brain pathological damages of AD patients.(Folke, Pakkenberg, & Brudek, 2019). Results of animal studies showed that blocking of the Wnt pathway could exacerbate the Tau hyper-phosphorylation, the formation and deposition of Aβ_1-42_, and the memory impairment (Tapia-Rojas & Inestrosa, 2018). Note that, abnormal function of Wnt signaling pathway can promote the production of Aβ, while Aβ, in turn, can inhibit the Wnt/β-catenin signaling pathway to aggravate the course of AD (Liu et al., 2014; Magdesian et al., 2008; Rodriguez et al., 2011).

It is a crucial experimental means and premise to establish effective animal models mimicking the pathophysiological process of human diseases. The main purpose of this study was to characterize the AD mouse model via bilaterally hippocampal microinjection of streptozotocin (STZ), a glucosamine-nitrosourea and DNA alkylating reagent. The behavioral performance was observed via the rotarod test (RT), open field test (OFT), tail suspension test (TST), sucrose preference test (SPT), novel object recognition (NOR) test, contextual fear conditioning (CFC) test, and Morris water maze (MWM). The serum concentrations of insulin and nesfatin-1 were tested via enzyme-linked immunosorbent assay (ELISA). The activation of microglia and astrocytes and neurogenesis in the hippocampus was detected by immunofluorescent staining with ionized calcium-binding adapter molecule 1 (Iba-1), CD68, glial fibrillary acidic protein (GFAP), and doublecortin (DCX) respectively. The expression of synaptic plasticity related molecules and key proteins of the Wnt pathway in the hippocampus and the prefrontal cortex (PFC) were detected via western blot.

## Results

### Bilaterally hippocampal injection of STZ induced a decreased tendency of body weight-gain

Figure 1 shows the bodyweight and bodyweight gain of mice in the two groups. As compared with that in the control group, the tendency of the bodyweight gain in the bilaterally hippocampal STZ-injected group was decreased, with a significant difference between groups during the second week post-operation.

**Figure 1:**
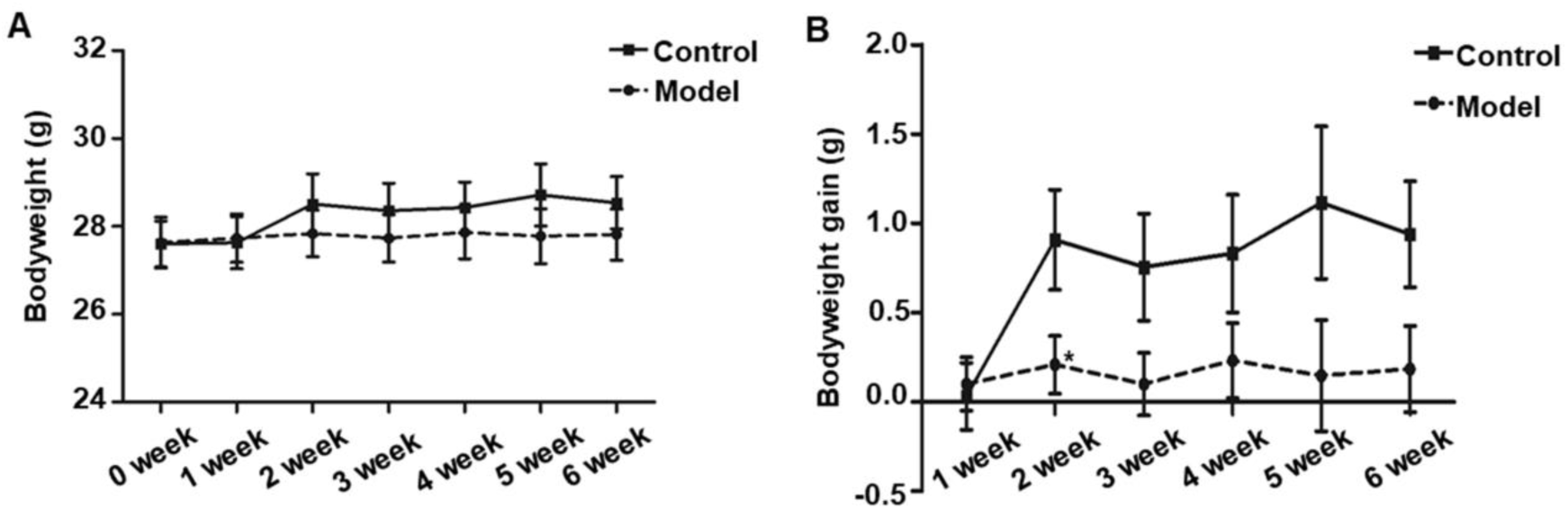
Effects of bilaterally hippocampal injection of STZ on the bodyweight gain in mice. The data are presented as the mean ± SEM, with 13 mice in each group. Although there was no significant difference between groups with regard to the bodyweight (A), the net bodyweight gain (B) of mice injected by STZ in the model group was lower than that in the control group in the second week. \**P*<0.05, \*\**P*<0.01; compared with the control group.

### Bilaterally hippocampal injection of STZ induced a decrease of locomotor activity and exploration behavior in mice in the OFT without significant changes in the RT, TST and SPT

General behavior performances of mice are shown in Figure 2, including the balance and motor coordination, automatic activity, exploratory and despair behavior, and anhedonia. As shown in Figure 2A, there was no significant difference between the two groups as regard to the falling time and falling speed in the RT. In the OFT, although there was no significant difference between groups with regards to the latency (Figure 2B) and frequency (Figure 2B) to the center, and duration in the center (Figure 2D), the bilaterally hippocampal STZ-injected mice showed significantly decreased total ambulatory distance traveling in the area (Figure 2C), as well as the ambulatory distance (Figure 2C) and ambulatory time (Figure 2D) in the center zone as compared with the control ones. The immobility time in the TST (Figure 2E) and the sucrose preference index in the SPT (Figure 2F) were not remarkably different between groups.

**Figure 2:**
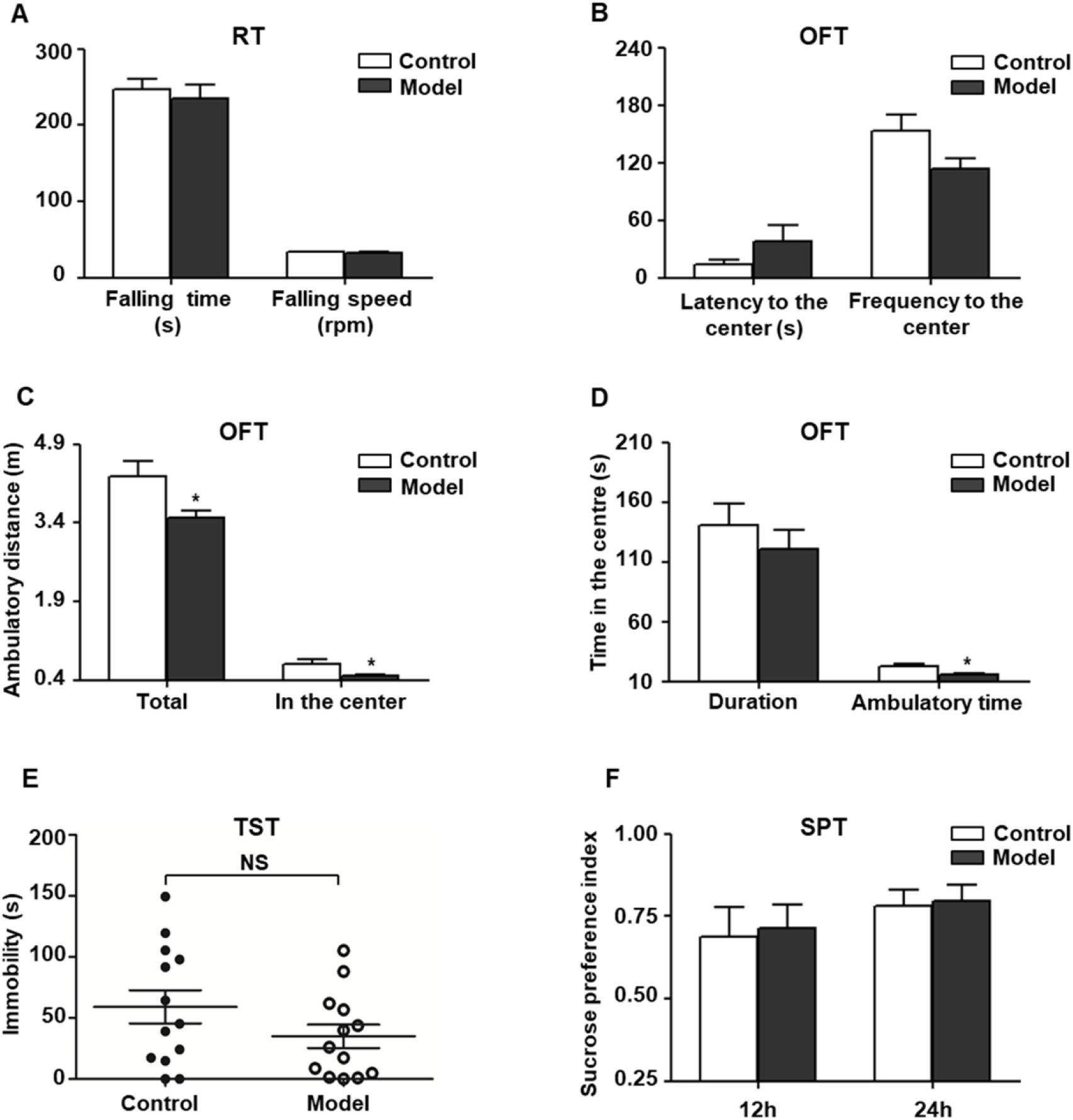
Effect of bilaterally hippocampal injection of STZ on the behavior performance of mice in the RT, OFT, TST and SPT. The data are presented as the mean ± SEM, with 13 mice in each group. There was no significant difference between the two groups as regard to the falling time (A) and falling speed (A) in the RT. In the OFT, although there was no significant difference in the latency (B) and frequency (B) to the center, duration (D) in the center between the two groups, bilaterally hippocampal STZ-injected mice showed significant decreased total ambulatory distance traveled (C) in the area, as well as the ambulatory distance (C) and time (D) in the center zone than control mice. The immobility time (E) in the TST and the SPI (F) in the SPT are not remarkably different between groups. \**P*<0.05, \*\**P*<0.01; compared with the control group.

### Bilaterally hippocampal injection of STZ induced an impairment of learning and memory ability in mice in the NOR and MWM, but not CFC

Behavioral performance of mice in the NOR task is shown in Figure 3A. As compared with the control mice, the bilaterally hippocampal STZ-injected mice spent less time exploring the novel object, with a significant decrease of the novel object recognition index.

**Figure 3:**
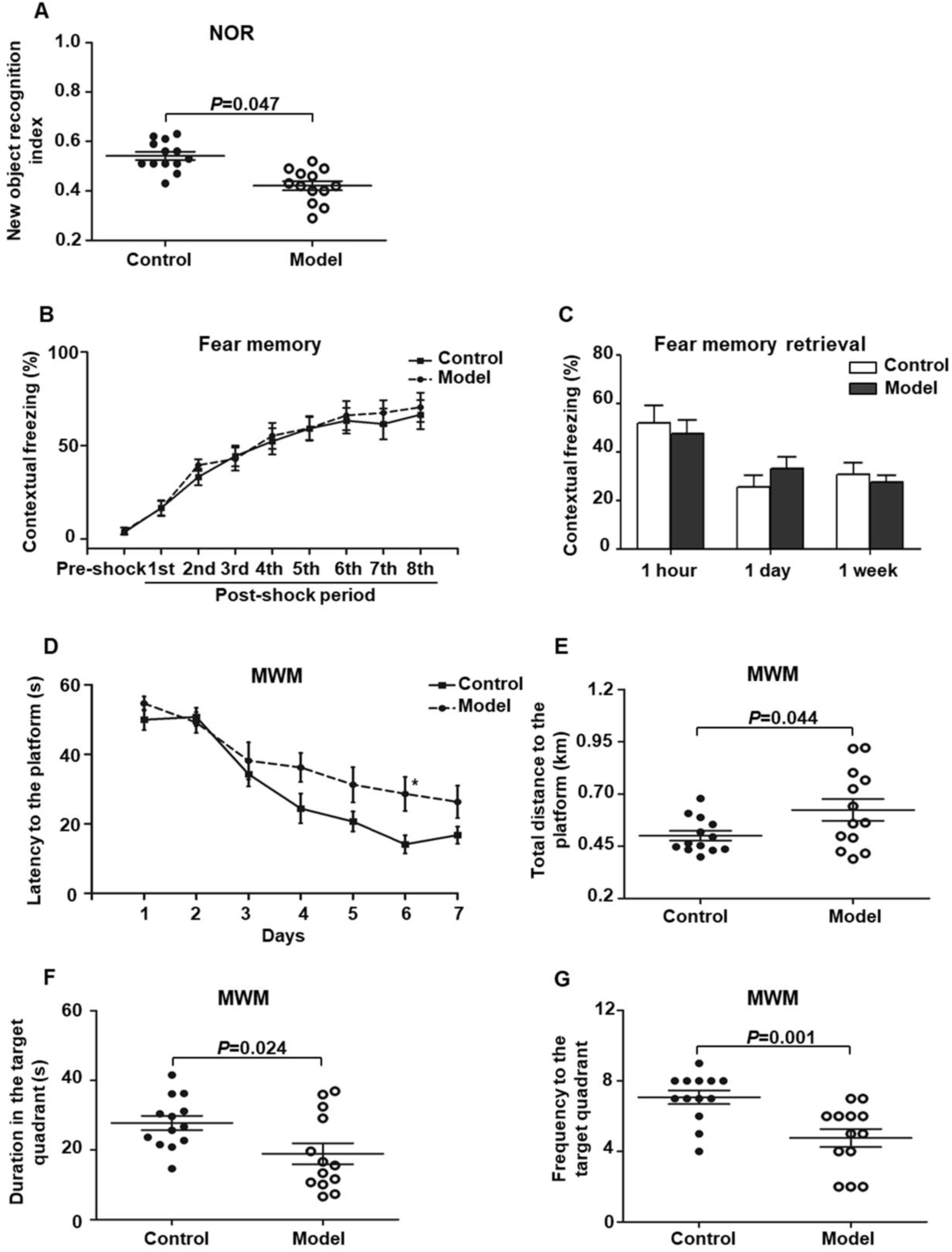
Effect of bilaterally hippocampal injection of STZ on the behavior performance of mice in the NOR, CFC and MWM. The data are presented as the mean ± SEM, with 13 mice in each group. In the NOR, compared with control mice, the novel object recognition index of model mice (A) was remarkably reduced. There was no significant difference between the two groups in the stage of fear memory formation (B) and freezing time of mice when the fear memory was extracted after 1 hour, 1day, and 1 week (C) in the CFC. In the MWM, during the acquisition phase, bilaterally hippocampal STZ-injected mice took longer time to find the submerged platform than the control mice did on sixth day (D). During the probe trial, the model mice showed increased total swimming distance to the platform (E), decreased duration (F) and less frequency (G) in the target quadrant. \**P*<0.05, \*\**P*<0.01; compared with the control group.

Figure 3B shows the performance of mice in the stage of fear memory formation in the CFC test, and there was no significant difference between the two groups.Moreover, the amount of freezing to context (Figure 3C) of the bilaterally hippocampal STZ-injected mice was not observably different from that of control ones, when the fear memories were extracted in the same environment 1 hour, 1 day and 7 days after formation of fear memory.

The performance of mice during the acquisition phase in the MWM is shown in Figure 3D. All the mice in this study learned how to locate the platform, as revealed by the sharp decline in the escape latency to the submerged platform. Moreover, the bilaterally hippocampal STZ-injected mice spent a longer time to find the escape platform than the control ones did on the sixth day. In the probe test, as compared with that of the control group, the total swimming distance to the platform (Figure 3E) of the model mice was significantly extended, together with a shorter duration (Figure 3F) in and less the frequency (Figure 3G) to the target quadrant.

### Bilaterally hippocampal injection of STZ increased the serum concentrations of insulin and nesfatin-1 in mice

As Figure 4 shown, compared with those in the control mice, the serum concentrations of insulin (Figure 4A) and nesfatin-1 (Figure 4B) were observably increased in the bilaterally hippocampal STZ-injected mice.

**Figure 4:**
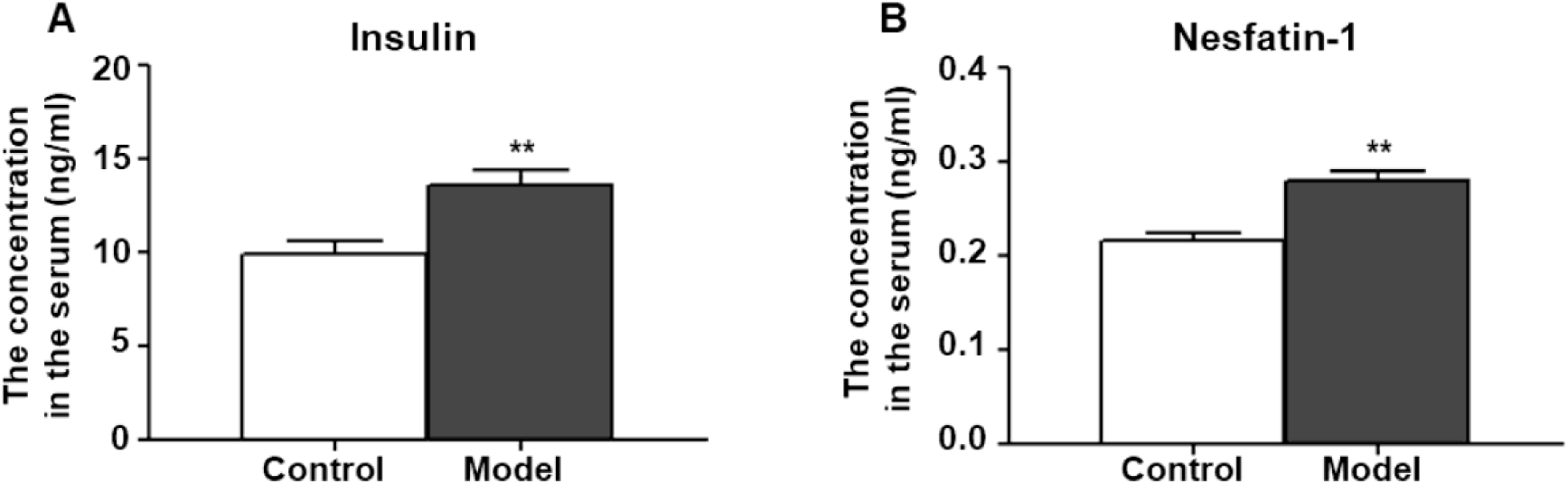
Effect of bilaterally hippocampal injection of STZ on the serum concentrations of insulin and nesfatin-1. The data are presented as the mean ± SEM, with 13 mice in each group. Compared with those in the control mice, the serum concentrations of insulin (A) and nesfatin-1 (B) were observably increased in the bilaterally hippocampal STZ-injected mice. \**P*<0.05, \*\**P*<0.01; compared with the control group.

### Bilaterally hippocampal injection of STZ promoted microglia to aggregate and activate in the hippocampus of mice

Figure 5A-F shows the typical immunofluorescent images of Iba-1 (green) and CD68 (red) in the mice hippocampal sections. As shown in Figure 5A and 5D, the Iba-1 (green) positive microglia clustered at the dentate gyrus (DG) of the bilaterally hippocampal STZ-injected mice, while such a phenomenon was not detected in the control group. Interestingly, as shown in Figure 5K, a dramatic ascending number of positive Iba-1 (green) marked microglia and CD68 (red) marked activated microglia were presented in the DG of the hippocampal tissues of model mice, and the activated microglial marker CD68 was mostly clustered around the hippocampal injection site. Figure 5G-J shows the typical immunofluorescent images of DCX (red) and GFAP (green) in the mice brain sections. There was no significant difference between groups as regard to the number of DCX (red) positive cells (Figure 5G, H and L). As shown in the Figure 5I, J and L, the GFAP (green) positive cells in the mice brain of the two groups were uniformly distributed in the hippocampus.

**Figure 5:**
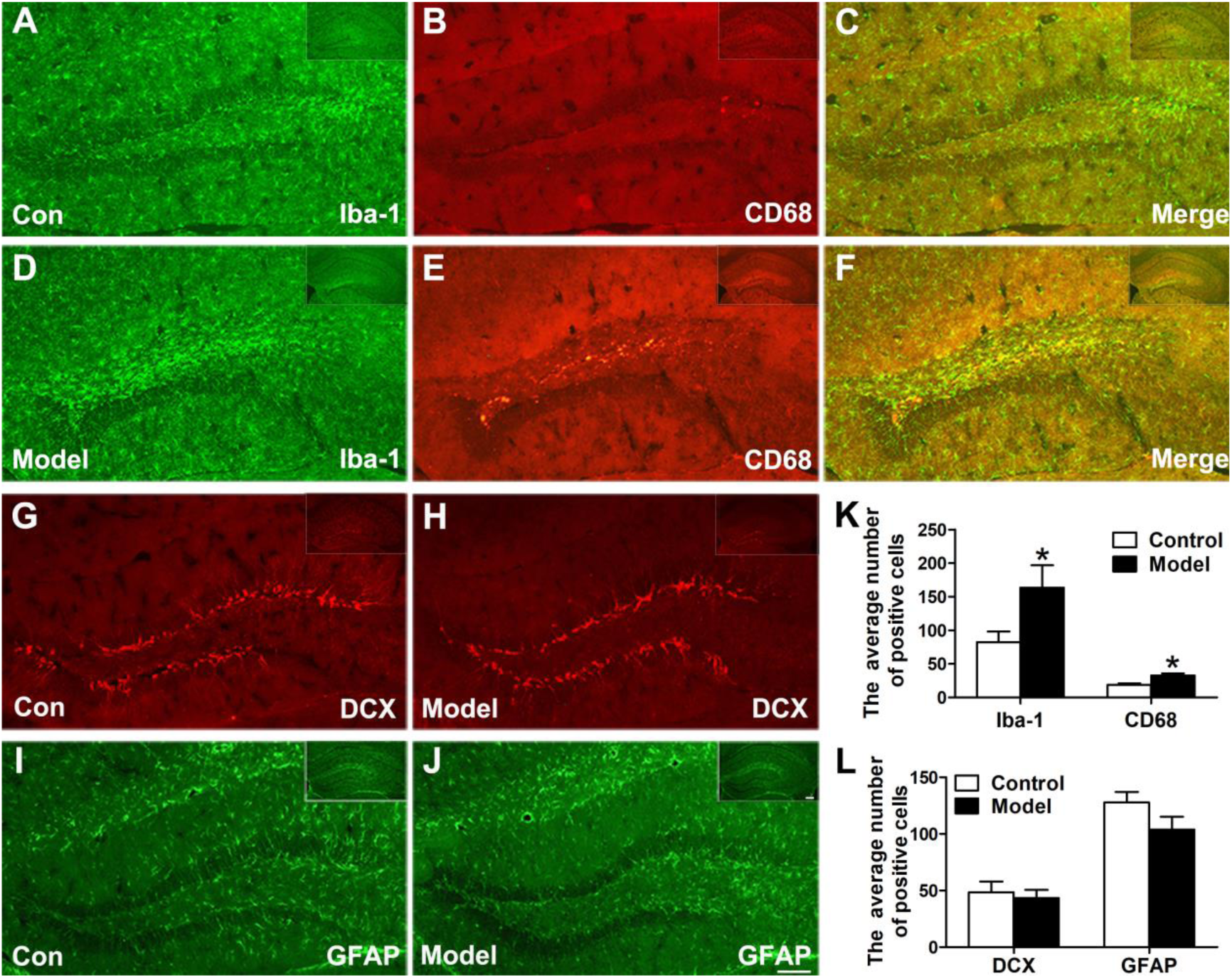
Effects of bilaterally hippocampal injection of STZ on the activation of microglias in the dentate gyrus of hippocampus of mice. A-F shows immunofluorescent microscopy of Iba-1 (green) and CD68 (red) in hippocampus tissues and the dentate gyrus. G-J shows immunofluorescent microscopy of DCX (red) and GFAP (green) in hippocampus tissues and the dentate gyrus (DG). Compared with those in the control mice, the number of Iba-1 (A, D) and CD68 (B, E) positive microglia ascends significantly in the DG tissues of the model group (K). There was no significant difference in the number of positive cells of DCX (G, H) and GFAP (I, J) between the two groups. The data in (K) and (L) are presented as the mean ± SEM, with 3 mice in each group. Scale bars = 100 μm in J (applies to A–J), 200 μm in insets. \**P*<0.05, \*\**P*<0.01; compared with the control group.

### Bilaterally hippocampal injection of STZ induced an imbalanced expression of synaptic plasticity related proteins in the hippocampus and the PFC in mice

Figure 6 shows the protein expressions of NMDAR2A, NMDAR2B, SynGAP, PSD95, and BDNF in the hippocampus and PFC of mice. Compared with those in the control mice, the expression levels of NMDAR2A, NMDAR2B, SynGAP, PSD95 and BDNF were remarkably decreased in both the hippocampus and the PFC of bilaterally hippocampal STZ-injected mice. Despite the small sample size, significant positive correlations were observed between the protein expression of BDNF and NMDAR2A (Hippocampus: r = 0.996, *P*<0.001, Figure S1A; PFC: r = 0.960, *P* = 0.002, Figure S1B), SynGAP (Hippocampus: r = 0.850, *P*=0.032, Figure S1C; PFC: r = 0.880, *P* = 0.021, Figure S1D), or PSD95 (Hippocampus: r = 0.857, *P* = 0.029, Figure S1E; PFC: r = 0.894, *P* = 0.016, Figure S1F).Moreover, significant positively correlations were also observed between the protein expression of NMDAR2A and NMDAR2B (Hippocampus: r = 0.820, *P* = 0.045, Figure S2A; PFC: r = 0.985, *P*<0.001, Figure S2B), SynGAP (Hippocampus: r = 0.848, *P* = 0.033, Figure S2C; PFC: r = 0.952, *P* = 0.003, Figure S2D), or PSD95 (Hippocampus: r = 0.851, *P* = 0.032, Figure S2E; PFC: r = 0.871, *P* = 0.024, Figure S2F).

**Figure 6:**
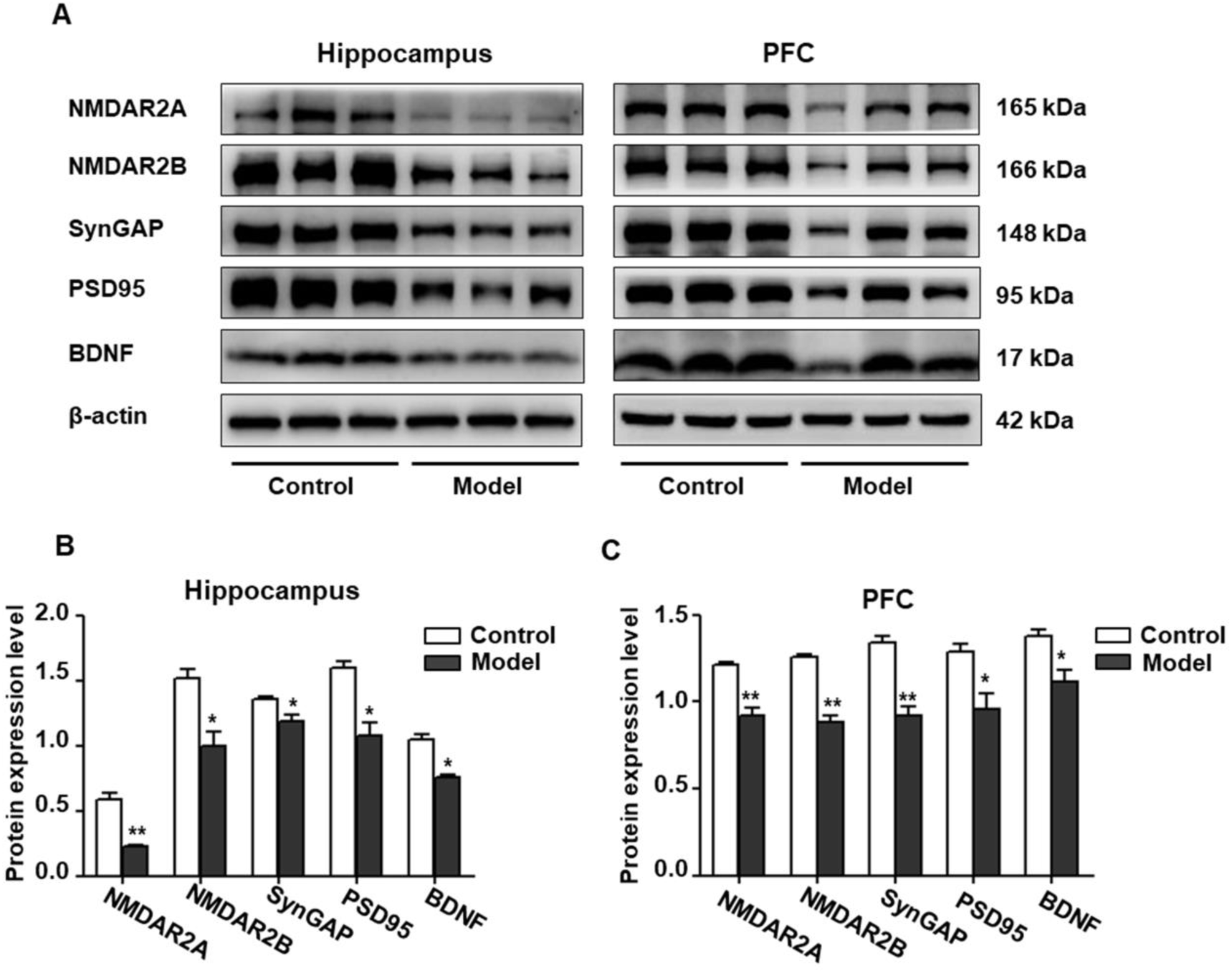
Effects of bilaterally hippocampal injection of STZ on the expression of synaptic plasticity related proteins in the hippocampus and PFC of model mice. Typical graph (A) and statistical analyses (B, C) of the western blot results. Data in (B, C) are presented as the mean ± SEM (n = 3 for each group). In the hippocampus (A, B), the expression of NMDAR2A, NMDAR2B, SynGAP, PSD95 and BDNF was all decreased in the model group. The proteins expression situation in PFC is similar to that in hippocampus (A, C). \**P*<0.05, \*\**P*<0.01; compared with the control group.

### Bilaterally hippocampal injection of STZ induced a dysregulation of Wnt/β-catenin signaling pathway in the hippocampus and the PFC in mice

Figure 7 shows the expression of key proteins in the Wnt/β-catenin pathway in the hippocampus and PFC of the mice. As compared with that in the control group, the relative expression of p-β-catenin/β-catenin in both the hippocampus and the PFC were remarkably declined in bilaterally hippocampal STZ-injected mice, and the expression of p-GSK-3β (ser9)/GSK-3β was decreased in the hippocampus, but not the PFC. No significant changes were found between groups as regard to the protein expression of CyclinD1 in both the hippocampus and the PFC. The results of Pearson’s correlation analysis showed that the protein expression of p-GSK-3β/GSK-3β was positively correlated with BDNF (r = 0.939, *P* = 0.005, Figure S3A), NMDAR2A (r =0.937, *P* = 0.006, Figure S3B), or synGAP (r =0.960, *P* = 0.002, Figure S3C) in the hippocampus.

**Figure 7:**
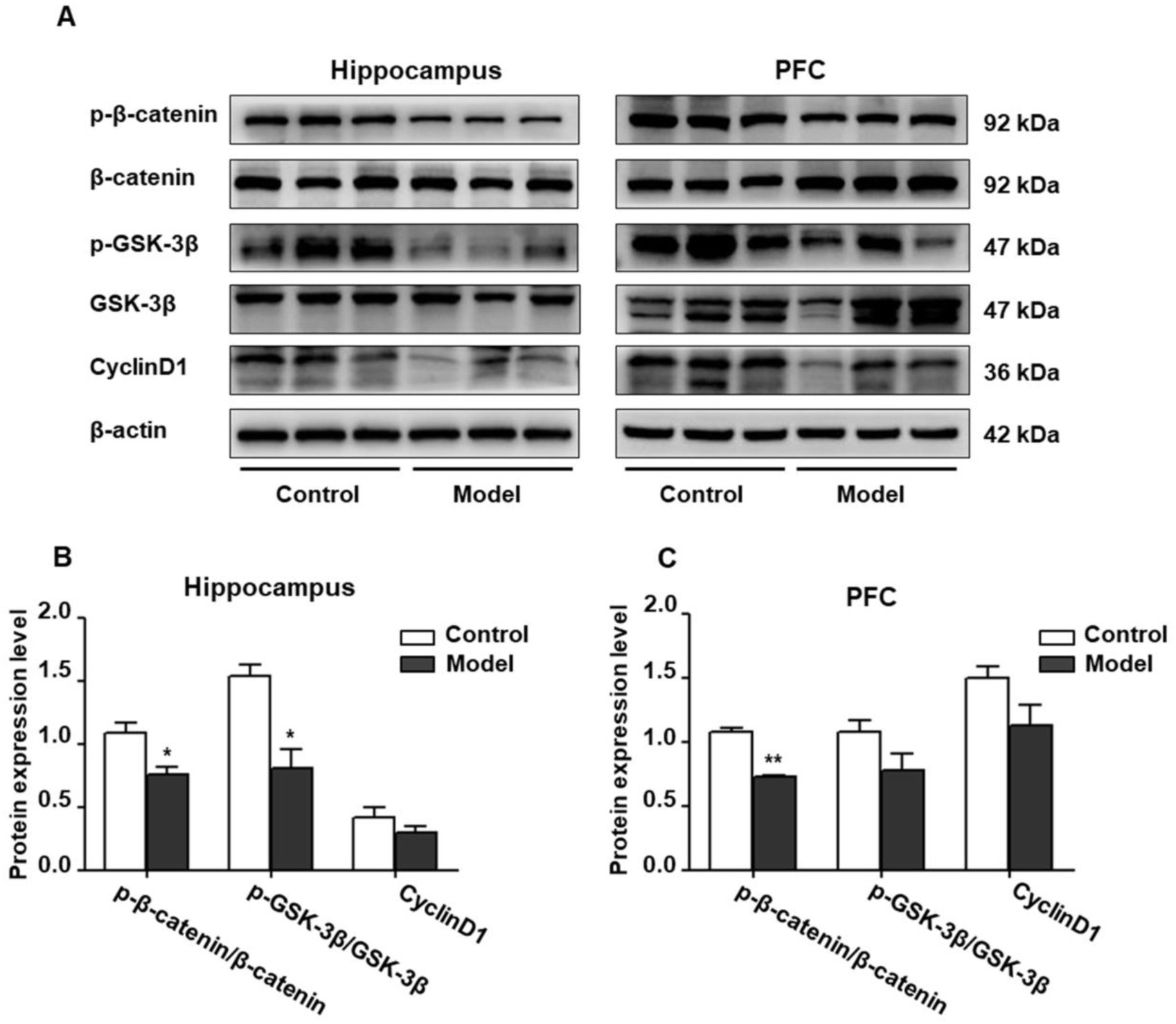
Effect of bilaterally hippocampal injection of STZ on the key protein expression of Wnt/β-catenin pathway in the hippocampus and prefrontal cortex of model mice. Typical graph (A) and statistical analyses (B, C) of the western blot results. Data in (B, C) are presented as the mean ± SEM (n = 3 for each group). In the hippocampus (A, B), the expression of p-β-catenin/β-catenin and p-GSK-3β(ser9)/GSK-3β was decreased in the model group, without CyclinD1. In PFC (A, C), although the expression of p-β-catenin/β-catenin of model mice was lesser than that of control mice, but not p-GSK-3β (ser9)/GSK-3β and CyclinD1. \**P*<0.05, \*\**P*<0.01; compared with the control group.

## Discussion

In the present study, an AD mouse model was established via bilaterally hippocampal injection of STZ. The results showed that a bilaterally hippocampal injection of STZ could induce a decreased tendency of bodyweight gain in mice, together with remarkably increased serum concentrations of insulin and nesfatin-1. Results of behavioral tests demonstrated that the spontaneous movement and exploratory behavior were decreased in the bilaterally hippocampal STZ-injected mice as compared with the control ones. More importantly, the model mice showed an task-specific impairment of learning and memory, as indicated by the declined novel object recognition index in the NOR, and the longer escape latency in the acquisition of MWM, along with the increased total swimming distance to the platform, decreased duration and less frequency in the target quadrant during the navigation phase of MWW. Moreover, the expression of NMDAR2A, NMDAR2B, SynGAP, PSD95, BDNF and p-β-catenin/β-catenin were remarkably reduced in the hippocampus and the PFC of the model mice, while the relative expression of p-GSK-3β/ GSK-3β was declined in the hippocampus. Furthermore, our results showed that the microglia in hippocampus of AD model mice was aggregated around the injection site, and the number of activated microglia increased remarkably in the DG. The alterations in the neuroendocrine response, the factors of synapse plasticity, and the parallel decline of learning and memory ability in the bilaterally hippocampal STZ-injected mice are in accordance with the network hypothesis of AD which proposes that compromised functionality of relevant neural networks may underlie the development of AD symptomatology.

STZ belongs to the family of glucosamine-nitrosourea compound and DNA alkylating reagent, and could be transferred to the cells alone by means of glucose transporters protein 2 (GLUT2). Peripheral injection of STZ is used widely to induce animal models of diabetes mellitus, as it could kill the pancreatic β cells high selectively at a dose-dependently manner and induce glucose metabolism disorder. Given the important role of insulin in the regulation of synaptic plasticity and the close relationship between diabetes mellitus and AD, STZ is also reported to be used in establishing AD animal models (Agrawal, Tyagi, Shukla, & Nath, 2011; Rai, Kamat, Nath, & Shukla, 2014). It has been reported that intracerebroventricular (icv) injection of STZ could cause not only the dysfunction of insulin/ insulin-like growth factor (IGF) pathway, but also the impaired learning and memory ability in mice (Agrawal et al., 2011). Consistently, in the present study, our results showed that bilateral injection of STZ into the dorsal hippocampus could induce a significant increase of serum insulin level in mice.

To investigate the behavioral changes induced by central administration of STZ and its subsequent increase of serum insulin concentration, a series of behavioral tests were performed in the present study. The results showed that hippocampal injected with STZ could induce an impairment of learning and memory in mice, together with decreased spontaneous movement and exploratory behaviors, which was consistent with the findings in previous studies of STZ icv injected mice (Moreira-Silva et al., 2018; Sorial & El Sayed, 2017; C. T. Zhang et al., 2016). There was no significant difference between groups as regard to the behavioral performance in the TST, SPT, and RT, indicating that a bilaterally hippocampal injection of STZ might not enough to induce despair, anhedonia, and impairments of balance and coordination activity in mice. On the whole, these results showed that bilateral hippocampal STZ injection could successfully induce a turbulence of insulin abundance and trigger task-specific AD-like behavior changes in mice, suggesting again the validity and reliability of STZ for establishing AD model. Moreover, considering the important role of hippocampus in maintaining the brain function, and the significant morphological atrophy and dysfunction of hippocampus in AD (Cavedo et al., 2014; Teixeira et al., 2018), hippocampal injection of agents has been attempted in exploring the pathogenesis of AD and evaluating the therapeutic effect (Karimi-Sales et al., 2020; Miao et al., 2019). In line with this, STZ was injected hippocampal microinjected bilaterally in this study and the injection site was verified by Nissl staining. Together with the reliable results in the behavioral tests, our results indicated that bilaterally hippocampal injection of STZ should be an effective method to establish an AD mice model, with the advantages of clear location, accurate dose and direct effect.

Nesfatin-1 is a satiety factor widely expressed in both central and peripheral tissues. Apart from the function of regulating feeding behavior and energy metabolism, nesfatin-1 has been reported to participate in the pathogenesis of anxiety, depression and other neuropsychiatric behaviors. Results of our previous study demonstrated that peripheral abundance of nesfatin-1 could be changed by high-fat diet and chronic stress, and ip injection of nesfatin-1 could induce anxiety and depression-like behavioral disorder in rats, together with the change of hippocampal synaptic plasticity (X.-X. Chen et al., 2019; Ge et al., 2015). Higher plasma level of nesfatin-1 was also reported in AD or diabetes mellitus patients and animal models (Alpua & Kisa, 2019) (Dong et al., 2013), with significant correlation to the cognitive dysfunction. In the present study, our results showed that the serum nesfatin-1 concentration was increased in the bilaterally hippocampal STZ-injected mice.Although the mechanism remains unclear, this results provide new evidence that nesfatin-1 might be involved with the insulin metabolism disorder-associated cognitive impairment in AD.

Neuro-inflammation has long been taken as a critical factor in the pathological of AD. It has been demonstrated that microglia can be activated by Aβ deposition, thereby secreting various inflammatory cytokines and inducing the activation of the complement system (Marttinen et al., 2018). In turn, the reactive inflammatory response will induce a further activation of microglia, as a vicious circle, ultimately leading to the neuronal dysfunction. The GFAP in the brain is a reliable manifestation of the astrocytes activation, Iba-1 is a specific marker of microglia, and CD68 is also used to label activated microglia. Increased expressions of Iba-1 and GFAP have been identified in the hippocampus of adult rats or mice received icv administration of STZ (Dos Santos, Vizuete, & Goncalves, 2020). In the present study, as compared with that of the control group, the positive cells labeled by Iba-1 and CD68 in the model group were aggregated in the dentate gyrus of hippocampal tissues, and the number of activated microglia cells was also significantly increased. The data is consistent with the phenomenon of a marked increase in CD68-labeled activated microglia in AD brain slices of APP/PS1 transgenic mouse (Gallagher et al., 2012). By contrast, there was no significant difference in GFAP positive expression of the STZ model group.

The doublecortin (DCX) is considered as a molecular marker of neurogenesis in hippocampal dentate gyrus and only expressed in the newborn neurons, and significantly reduced number of newborn neurons labeled with DCX was demonstrated in the hippocampus of an AD rat model induced by icv injection of STZ (Mishra, Singh, Shukla, & Shukla, 2018). In this present study, there was just a downward trend of DCX immune-positive cells in the dentate gyrus of the model mice, without a significant difference between groups. These results indicated that the reactive inflammatory response was activated after bilaterally hippocampal injection of STZ, together with subtle decline of neurogenesis. These changes might interpret partly the impairment of learning and memory of AD mice induced by STZ hippocampal injection. However, more detailed investigation should be carried out focusing on the delicate shifts and the potential mechanism in the future.

Brain-derived neurotrophic factor (BDNF) can provide neurotrophic support for different neuronal populations including 5-HT and is a seminal regulator of synaptic plasticity. Decreased BDNF concentration has been reported in serum and CSF of AD patients, with a significantly positive relationship between the degree of decline and the severity of cognitive dysfunction (Li et al., 2009; Ng, Ho, Tam, Kua, & Ho, 2019). Moreover, it has been demonstrated that lateral ventricle injection of nerve growth factor can significantly improve the cognitive abilities of AD transgenic mice by up-regulating the hippocampal expression of BDNF. Furthermore, BDNF can reverse neuron loss, improve cognitive impairment, and provide substantial protection for important neural circuits associated with AD (Nigam et al., 2017). Consistent with these findings, our results showed a decreased protein expression of BDNF in the hippocampus and the PFC of bilaterally hippocampal STZ-injected mice, suggesting the central role of BDNF in the pathogenesis of AD again. It has been reported that astrocytes could secrete BDNF to promote the development and survival of the central nervous system (de Pins et al., 2019). However, in the present study, the results of immunofluorescent staining showed no significant difference between groups about the number of astrocytes in the DG, indicating that the decreased expression of BDNF in the hippocampus and PFC of STZ-injected mice may not be regulated totally by the astrocytes.

NMDARs serve as modulators of synaptic transmission in the mammalian central nervous system with both short-term and long-lasting effects, and take responsible for maintaining the neuronal excitability, Ca^2+^ influx, and memory formation (Kamat et al., 2016). BDNF is reported to bind with TrkB and promote the activation of NMDAR and the aggregation of PSD95 through the PI3K-Akt pathway to regulate synaptic plasticity (Yoshii & Constantine-Paton, 2007). Over-activation of NMDAR could lead to neuronal cell death, excessive influx of Ca^2+^, and abnormal generation of free radicals, which drives synaptic dysfunction and hyper-phosphorylation of Tau protein (Rai, Kamat, Nath, & Shukla, 2013). Memantine, the NMDA receptor antagonist, has been reported to improve the cognitive function of AD patients by improving the activity of glutamatergic neurons (Hong, Choi, Jeong, Park, & Na, 2018; Jason & Andrew, 2018). Besides, the content of phosphorylated and non-phosphorylated NMDAR subunits in the hippocampus of AD patients are reduced (Sze et al., 2001). Consistently, in SAMP8 mice or AD models injected with Aβ_25-35_ or Aβ_1-42_, the expression of NMDAR1, NMDAR2A and NMDAR2B were decreased markedly in the hippocampus or cortex (K. W. Chang et al., 2020; K. Wang et al., 2018; Xu et al., 2016). Moreover, it has been demonstrated that ketamine, a noncompetitive NMDAR antagonist, could ameliorate the impaired learning and memory of developing mice induced by repetitive mechanical stress, and the mechanism may be associated with the increased hippocampal BDNF expression (C. H. Chang, Su, & Gean, 2018; Peng, Zhang, Wang, Ren, & Zhang, 2011). In this present study, the expression of NMDAR2A and NMDAR2B in the hippocampus and PFC of mice were significantly down-regulated after bilaterally hippocampal injection of STZ, and a significant positive correlation was found between the protein expression of BDNF and NMDAR2A. These results further confirmed the central role of BDNF in the pathogenesis of AD and the close relationship between BDNF and NMDARs.

Synaptic plasticity is defined as the specific changes in the number, structure and function of synapses due to the continuous neuron activity, which is the material basis for regulating learning and memory functions and emotional states. Synaptic plasticity divides into synaptic transmission and synaptic structural plasticity, both of which are closely related to the pathogenesis of AD. The PDZ domain of PSD95 can be combined with the intracellular domain of the NMDAR2 subunit, thereby anchoring NMDAR to the post-synaptic densities, based on which NMDAR can exert a biological function on synapses (X. Chen et al., 2015). Unsurprising, PSD95 is reported decreased in the hippocampus of APP/PS1 AD transgenic mice (X. Zhang et al., 2020). SynGAP, a protein enriched in excitatory synapses, can integrate with PSD95, Shank and Ca^2+^/CaMKII to form a complex that plays a key role in synaptic plasticity (Kim, Liao, Lau, & Huganir, 1998). Aβ oligomers have been reported to be capable of instigating synapse dysfunction and deterioration via binding with SynGAP, subsequently aggravating the course of AD (Ding et al., 2019). Results of postmortem study showed that the SynGAP levels in the PSD were remarkably declined in the PFC of AD patients (Gong, Lippa, Zhu, Lin, & Rosso, 2009). Compared with that of control group, the expressions of SynGAP and PSD95 in the hippocampus and the PFC of AD model mice was significantly down-regulated in the present study. Apart from the significant positive correlation between BDNF and NMDAR2A, the protein expression of SynGAP and PSD95 were also remarkably correlated with the expression of BDNF. Together with the shifted expression of BDNF and NMDARs in this study, our result suggests again that the synaptic plasticity, involved with the balance of multiple proteins and multiple receptors, play a critical role in the pathogenesis of AD.

Dysfunction of the Wnt/β-catenin signaling pathway is also involved in the progress of AD, which is manifested by the decreased level in β-catenin and increased activity of GSK-3β. The inhibition of Wnt/β-catenin pathway is reported to promote massive accumulation of Aβ and Tau hyper-phosphorylation, and Aβ in turn inhibits the Wnt/β-catenin pathway, eventually resulting in neurofibrillary tangles in AD patients and worsening the course of AD (Jia, Pina-Crespo, & Li, 2019). Moreover, it has been demonstrated that blocking Wnt/β-catenin pathway could down-regulate the expression of BDNF induced by NMDAR activation in the primary cortical neurons of mice, whereas activation of Wnt/β-catenin pathway could stimulate and increase BDNF expression (W. Zhang et al., 2018). NMDAR was also found to regulate the expression of disintegrin and metalloproteinase 10 through the Wnt/MAPK signaling pathway, which is a novel target for the treatment of AD (Wan et al., 2012). In this present study, the phosphorylation of β-catenin and GSK-3β (ser9) were decreased in the hippocampus or PFC of mice after bilaterally hippocampal injection of STZ, although the expression of CyclinD1 was not significantly different between groups. The phosphorylation of GSK-3β at ser9 inhibited the activity of GSK-3β, and the reduced phosphorylation of GSK-3β (ser9) in the hippocampus indicated the abnormal increased activity of GSK-3β. It has been demonstrated that GSK-3β could directly inhibit the expression of BDNF by phosphorylation of CREB (Grimes & Jope, 2001). In the present study, the protein expression of p-GSK-3β/GSK-3β was positively correlated with BDNF. These results indicated again the expression pattern of Wnt/β-catenin signaling pathway might be associated with the pathogenesis of STZ-induced AD, and BDNF might be taken as an important factor linking the imbalanced protein expression to the changed synaptic plasticity.

There were several limitations in the current study. First, although the changes of behavioral performance and the central expression of BDNF and many molecular associated with synaptic plasticity were observed, the abundance of Aβ_42_ and Tau were not detected in the study. Second, although the central of BDNF in the AD-like neurobiological changes induced by bilateral hippocampal STZ injection was suggested, it was not further identified by intervening the abundance or function of BDNF. Moreover, the correlation between BDNF and insulin or nesfatin-1 need to be observed in our further studies. In summary, our results showed that a bilaterally hippocampal injection of STZ could successfully induce AD-like behaviors in mice. Moreover, dysfunctional neuroendocrine response was also found in the bilaterally hippocampal STZ-injected mice, as indicated by the increased serum concentration of insulin and nesfatin-1. Apart from the hippocampal inflammatory response, the mechanism might be associated with the imbalanced expression of synaptic plasticity-related proteins in hippocampus and PFC, including NMDAR2A, NMDAR2B, SynGAP, PSD95 and BDNF. Moreover, the dysfunction of wnt/β-catenin pathway, especially the decrease in the phosphorylation of β-catenin and GSK-3β, may also be involved in AD-like pathological process. Given the relationship between BDNF and the imbalanced expression of proteins associated with regulation of synaptic plasticity in the hippocampus and the PFC, the central role of BDNF was self-evident. These findings might provide new evidence for understanding the pathogenesis of STZ-associated AD.

## Methods and Materials

### Animals

Twenty-six adult C57BL /6 male mice (bodyweight 25 ∼ 30 g, aging 3 months) were purchased from and kept in the experimental animal center of Tongji University School of Medicine under the condition equipped with a commutative 12 h light/dark schedule. The mice were randomly divided into a control group (CON) and a STZ-injection model group (MOD) with 4-5 mice per cage and access to food and water *ad libitum*. The ambient temperature was maintained at 20 ± 2 °C with 50 ±5% relative humidity. All the animal care practices and experiment procedures were reviewed and approved by the Animal Experimentation Ethics Committee of Tongji University School of Medicine.

### Stereotactic intra-hippocampal microinjection of STZ and experiment design

Each mouse was anesthetized by intraperitoneal injection of 5% chloral hydrate (0.1 ml / 10 g), secured to a platform placed in a stereotaxic apparatus (David Kopf; Tujunga, CA, USA) and the scalp incised. After the bregma was located, a rotary tool was used to drill into the skull and get two burr holes, according to the following sites of 2.1 mm posterior, 1.3 mm lateral and 1.8 mm below. Mice in the MOD groupreceived bilateral infusions of STZ (Sigma-Aldrich, s0130; 3 mg/kg) dissolved in 0.9% sterile saline into the hippocampus on both coordinates with a constant rate (500 nL/min) pulled by a micropipette puller. Correspondingly, mice in the CON group received bilateral infusions of 0.9% sterile saline. After the wound was closed, the animals were placed on the 37°C warming pad to recover until their respiration and locomotor activity returned to normal (C. T. Zhang et al., 2016).

Bodyweight of mouse was monitored once a week. After 2 weeks of recovery from surgery, all mice were submitted to a chain of behavioral paradigms. The schedule of the experimental design is shown in Figure 8.

**Figure 8:**
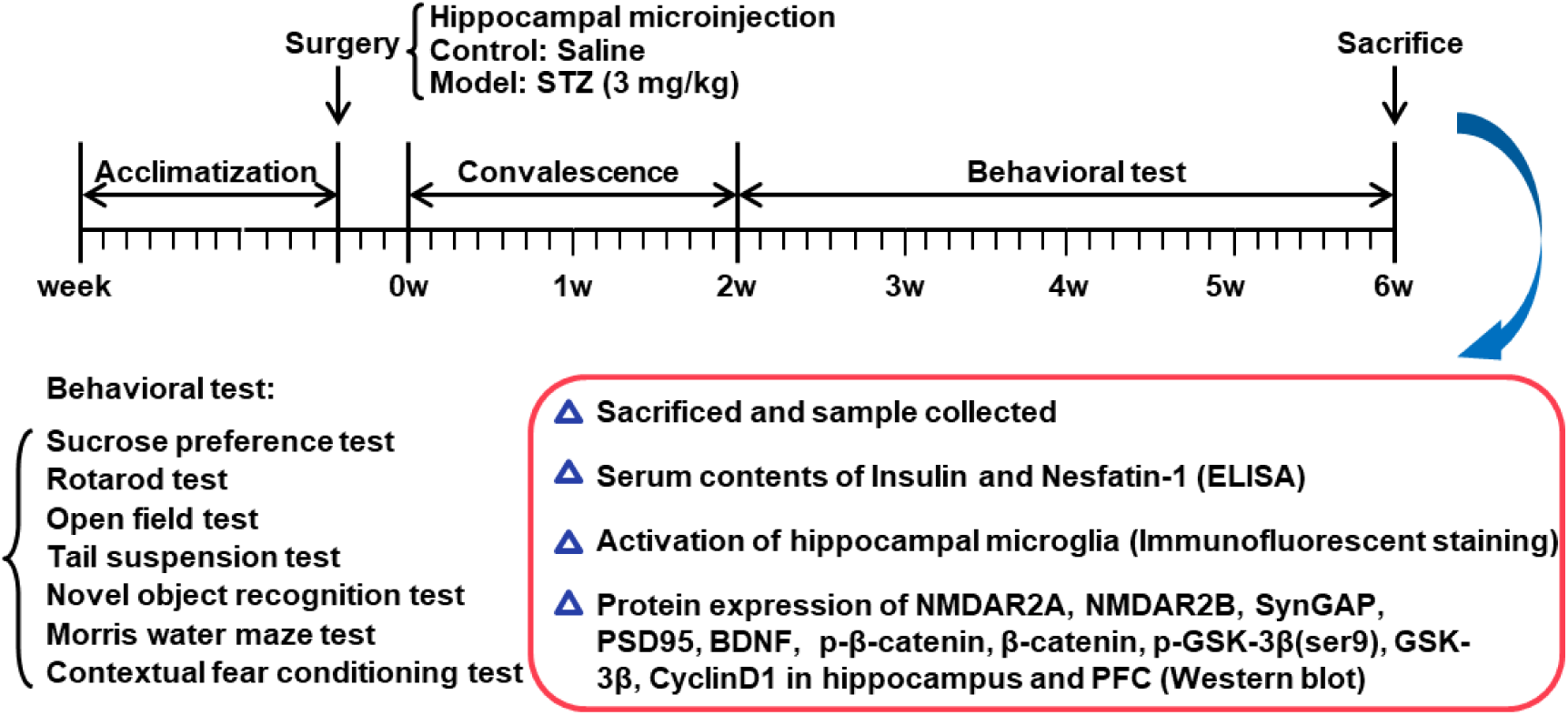
Schedule of the experimental design. STZ, streptozotocin; BDNF, brain derived neurotrophic factor; PFC, prefrontal cortex.

### Behavioral test

All behavioral experiments were performed during the light phase of the light/dark cycle and in a sound-proof room with a neutral environment. All the mice in this study were given a 30-min habituation time after transported to the experimental rooms and subjected to the following behavioral tests. The experimenter was blind to the group identity of the tested mice. The behavioral tests were recorded with a camera and a trained researcher analyzed these videos.

#### Sucrose preference test (SPT)

The SPT was used to detect the anhedonia behavior in rodents (Ke et al., 2020). After a deprivation of food and water for 12 hours, the mice were housed solely and had free access to two identical water bottles filled with either sterile water or 1% (w/v) sucrose solution respectively. To eliminate the interference of the location factor, the positions of the bottles were swapped after 12 hours. The mass of water and sucrose solution consumed were weighed after 12 hours and 24 hours respectively. The sucrose preference index (SPI) is defined as the ratio of the sucrose water ingested to the total amount of liquid intake.

#### Rotarod test (RT)

Motor coordination and balance of mice was assessed by RT in a reliable way. The test was divided into two stages, namely the training period and the test period. The training period consisted of a uniform 10 rpm for 1 min, and an acceleration period of 4-40 rpm for 5 min with relaxing 30 min at least. The mice were placed on the rotating apparatus with their backs to the operator. Note that, once the mouse fell during the training, it will be put back immediately to make sure completion. The test period was conducted on the second day after the training by three trails on the rotating rod, and inter-trial interval was 1 h for each mouse. During the process of accelerating the rod device from 4-40 rpm over the course 5 min, the latency to fall off and the maximal rotation rate before the mouse fell down were recorded and evaluated (Kandimalla, Manczak, Yin, Wang, & Reddy, 2018).

#### Open field test (OFT)

The OFT was conducted to synchronously explore the locomotion and exploration. The apparatus was made up of a square arena, with a white floor divided into 9 squares (10 cm × 10 cm) and enclosed by continuous 20.3 cm-high walls made of transparent plexiglass. Each mouse was put into it at one corner and headed toward the wall. The total distance traveled, latency to the central square, distance and duration in the central square, as well as the frequency through the central square over a 30 min period were recorded and analyzed by Activity Monitor software (Med Associates, St. Albans, VT, United States) (Q. Zhang, Huang, Zhang, Ding, & Song, 2019).

#### Tail suspension test (TST)

The TST was conducted as previously described (Zhou et al., 2016) with some modifications. Briefly, mice were individually suspended by tail (20 cm from floor) using a paper tape (1.5 cm from the tip of the end). Each mouse was suspended for a total of 6 min, and the duration of immobility was recorded during the final 4 min interval of the test. Mice were considered immobile only when they hung passively and completely motionless. In addition, the mouse that once climbed up their tails was not involved in this assay.

#### Novel object recognition (NOR) test

The NOR test was performed to assess non-spatial memory in a 25 cm × 25 cm × 25 cm and black soundproof box (Zhu et al., 2014). The test consisted of three phase: habituation, training, and retention sessions. In the first stage, the single mouse was placed into the box facing the wall, having a preceding 10 min habituation once a day, for 3 consecutive days. During the training session, two indistinguishable objects were symmetrically fixed to the floor of the box, and each mouse was allowed to travel in the apparatus freely for 10 min. After the training, the mice were immediately returned to their home cages for 60 min at least. Ultimately, during the retention test, each mouse was placed back into the same box, with one of the familiar objects substituted by a novel object. Test mouse was then allowed to freely explore for 5 min and videotaped. New object recognition index, a ratio of the amount of time spent exploring the novel object over the total time spent exploring the both, was used to evaluate cognitive function. A mouse was scored as exploring the object when touching or sniffing the object with its nose or front legs, and facing the object at a distance within 1 cm (D. Wang, Wang, Liu, & Li, 2018).

#### Morris water maze (MWM) test

MWM serves as a method of evaluating different aspects of spatial memory in rodents. The maze was equipped with a circular blue pool (diameter 120 cm, height 60 cm, filled with water to 40 cm high at 21°C), which was divided into four hypothetical, equal quadrants. A circular platform (11 cm diameter) was submerged approximately 1 cm below the surface of water in the center of the target quadrant. Several complex visual cues surrounded the pool diagonally and were used for facilitating orientation. The test consisted of a 7-day acquisition phase and a 1-day probe test on the 8th day.During the acquisition phase, each mouse received 4 trials per day, and was trained to find the hidden platform. The mice was placed into the pool facing the wall individually to find the submerged platform within 60 seconds, and stayed on it for 20 s. If the hidden-platform was not found within 60 s, the mouse was gently guided to the platform and held for 20 s. On the 8th day, the probe test was performed to determine memory retention, and the platform was removed. Each mouse was put into the pool from the diagonally opposite side of the previous platform, and permitted to swim for 60 s freely. The escape latency in the acquisition phase, total distance to the platform, the duration spent in and frequency to the target quadrant in the probe test were monitored and measured automatically by Noldus software (EthoVision XT 8.0, Noldus Technology) (Kandimalla et al., 2018).

#### Contextual fear conditioning (CFC) test

The CFC paradigm was considered to assess the ability of associating a distinct context with aversive footshocks through hippocampus-dependent cognitive mechanisms in rodents as described previously (Dai et al., 2008). The mouse was subjected to a CFC test in an sound attenuating chamber (30 × 25 × 20 cm), the floor of which was consisted of parallel 2 mm diameter stainless steel rods spaced 8 mm apart. A preceding 10 min habituation was subjected to reduce the contribution of anxiety and stress on the outcome. 24 h later, mice were placed in the same chamber and allowed to explore freely for 120s before receiving 8 unsignaled foot shocks (1.0 mA, 2 s) with inter-shock intervals of 60s. Mice were immediately returned to their home cages 2 min after the final foot shock. The behavioral freezing to the context of mice was measured to evaluate contextual fear memory at three points: 1 h, 24 h, and 1 week after fear conditioning. The mice were placed back to the same context and scored freezing behavior for 660s by the automated FreezeFrame system (Coulbourn Instruments), which digitizes the video signal at 4 Hz and compares movement frame by frame to score the amount of freezing. Freezing was defined as the absence of all movement except for that needed for breathing (Devi & Ohno, 2013).

### Measurement of the serum concentrations of insulin and nesfatin-1

Twenty-four hours after the last behavioral test, all mice were sacrificed and the blood was collected. The serum levels of insulin and nesfatin-1 were detected using commercially available enzyme-linked immunosorbent assay (ELISA) kits (insulin: Yuanye Biotech. Co., LTD, Shanghai, China; nesfatin-1: Cusabio Biotech. Co., LTD, Wuhan, China) according to the manufacturers’ instructions.

### Immunofluorescent staining

Three mice in each group were selected randomly and transcardially perfused with 0.01 M phosphate-buffered saline (PBS, pH 7.4) followed by 4% paraformaldehyde (PFA) in PBS. The whole brains were dissected out and post-fixed in 4% PFA overnight, followed by cryoprotection in 30% sucrose in PBS overnight. After embedded with optimum cutting temperature compound (OCT), the brains were cut into sagittal sections (25 m thick), and then stored at -80°C for immunostaining (Pruski et al., 2019).

Immunohistochemistry was performed as described previously (Zhu et al., 2014) with some modifications. Sections were re-hydrated washed in PBS for 3×10 min in a microwave followed by incubation with a blocking solution containing 5 % bovine serum albumin (BSA) and 0.1 % Tween-20 in PBS (pH 7.4) for 6 min at 95 °C to retrieve the antigen. Then, brain sections were incubated with rabbit anti-Iba-1 (1:1000; LAK4357, Wako, Japan), rat anti-CD68 (1:200; MCA1957GA, Bio-Rad Company, United States), guinea pig anti-DCX (1:300; AB2253, Millipore, USA) or rabbit anti-GFAP (1:1000; z0334, Dako, USA), at 4°C overnight. After several washes in PBS, sections were incubated with donkey anti-rabbit Alexa Fluor 488-conjugated IgG (1:500; R37118, Invitrogen, Carlsbad, CA, USA), and biotinylated horse anti-rat or horse anti-guinea pig IgG (1:500, Jackson ImmunoResearch, USA) at room temperature for 2 h. Ultimately, brain sections were incubated with streptavidin-Cy3 (1:1000, 016160084, Jackson ImmunoResearch, West Grove, PA, USA) and counterstained with Hoechst 33258 (1:2000, 94403, Sigma, St.Louis, MO, USA) at room temperature for 1 h. Fluorescent images were taken on a laser-scanning confocal fluorescent microscope (Leica TCS SP8, Leica Microsystems, Wetzlar, Germany).

### Western blot assays

Another 3 mice were selected randomly in each group, and sacrificed by cervical dislocation 24 hours after the last behavioral test. The hippocampus and the PFC were dissected promptly, frozen in liquid nitrogen and stored at –80°C. The western blot protocol was carried out as described in our previous study. Briefly, the tissues were lysed in ice-cold radio immunoprecipitation assay (RIPA) buffer containing protease inhibitor cocktail (Roche, IN, USA) and the phosphatase inhibitor PhosSTOP (Roche, IN, USA). Determine equal amounts of proteins (20 µg) using the BCA Protein Assay Reagent Kit before loading onto the 12.5% SDS-PAGE. After separated from each other, the proteins were subsequently transferred onto the polyvinylidene difluoride membrane (PVDF). The membrane was blocked 2 h with 5% BSA in TBST at RT followed by incubation with primary antibodies respectively at 4 °C overnight. The primary antibodies were as follows: anti-NMDAR2A (1:2000; 19953-1-AP; Proteintech), anti-NMDAR2B (1:2000; 21920-1-AP; Proteintech), anti-SynGAP (1:2000; 19739-1-AP; Proteintech), anti-PSD95 (1:2000; MA1-045; Thermo), anti-BDNF (1:1000; sc-546; Santa Cruz), anti-p-β-catenin (1:1000; 9561; Cell Signaling Technology), anti-β-catenin (1:1000; 9562; Cell Signaling Technology), anti-p-GSK-3β(ser9) (1:1000; 9323; Cell Signaling Technology), anti-GSK-3β (1:1000; 9315; Cell Signaling Technology), anti-CyclinD1(1:1000; ab190564; Abcam) and β-actin (1:2000; TA-09; Zhongshan Biotechnology). Then the membranes were further incubated with a horseradish peroxidase-conjugated secondary antibody, goat anti-rabbit or goat anti-mouse secondary antibodies (Abcam), at RT for 1 h. All procedures were performed by rinsing in TBST for 10 min and repeated three times. After developed with the Easy Enhanced Chemiluminescence Western Blot Kit (Pierce Biotechnology, Rockford, IL, USA), the protein blots were visualized by a gel imaging system (Tanon Science& Technology Co., Ltd., Shanghai, China) and then images were processed and analyzed using Image J software (NIH) (X.-X. Chen et al., 2019).

### Statistical analysis

All statistical analyses were performed using SPSS (version 12.0.1, SPSS Inc., Chicago, IL, United States). The difference between groups was examined using Student’s *t*-test. Data are expressed as means ± SEM and *P*< 0.05 was considered statistically significant. Correlation analysis was performed using a Pearson’s correlation test.

## Abbreviations

Aβ: amyloid beta;
AD: Alzheimer’s disease;
ADDLs: Aβ-derived diffusible ligands;
CFC: contextual fear conditioning test;
CREB: cAMP response element-binding protein;
DCX: doublecortin;
ELISA: enzyme-linked immunosorbent assay;
FDA: American Food and Drug Administration;
GFAP: glial fibrillary acidic protein;
GLUT2: glucose transporters protein 2;
Iba-1: ionized calcium-binding adapter molecule 1;
IGF: insulin-like growth factor;
MWM: Morris water maze;
NFTs: nerve fiber tangles;
NMDARs: N-methyl-D-aspartate receptor;
NOR: novel object recognition test;
OFT: open field test;
PFC: prefrontal cortex;
PSD95: postsynaptic density protein 95;
RT: rotarod test;
SPs: senile plaques;
SPI: sucrose preference index;
SPT: sucrose preference test;
STZ: streptozotocin;
SynGAP: Synaptic Ras GTPase activation protein;
TST: tail suspension test.

## Acknowledgements

The authors would like to thank Dr. Qiong Zhang and Ling Hu who assisted in the experiments of molecular biology and statistical analysis of data.

## Role of the funding source

Funding for this study was provided by the Natural Science Foundation of China (81401122, 81870403). These institutions had no role in the study design; the collection, analysis, or interpretation of the data; the writing of the manuscript, or the decision to submit the paper for publication.

## Author contributions

C.C.Q and X.X.C performed most of the experiments, analyzed the data, and wrote the manuscript. X.R.G performed Western blotting analysis. J.X.X and S.L modified the pictures. C.C.Q and J.F.G participated in the design of the study. J.F.G conceived the study and revised the manuscript. All authors read and approved the final manuscript.

## Competing Interests

The authors have declared that they have no actual or potential competing interest exists associated with the conduct of this work.

## Notes

### Competing Interest Statement

The authors have declared no competing interest.

