## supplemental figures for "Impaired learning and memory ability induced by a bilaterally hippocampal injection of streptozotocin in mice: involved with the adaptive changes of synaptic plasticity"

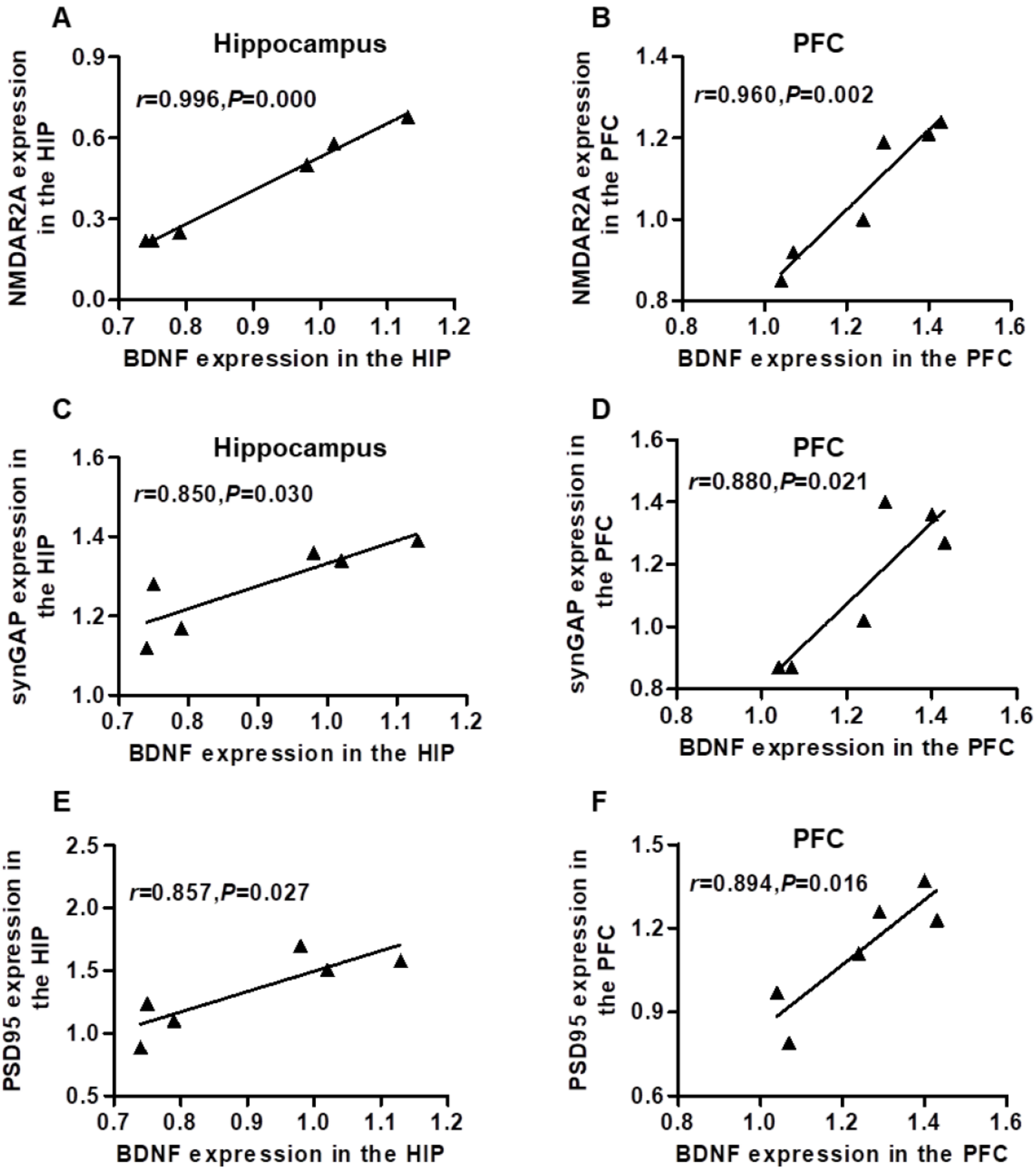


**Supplement 1:** Correlation analysis of the protein expressions between BDNF and NMDAR2A, synGAP, PSD95 in the hippocampus and PFC.

In the hippocampus and PFC, BDNF expression was positively related to the expression of NMDAR2A (A, B), synGAP (C, D) and PSD95 (E, F).

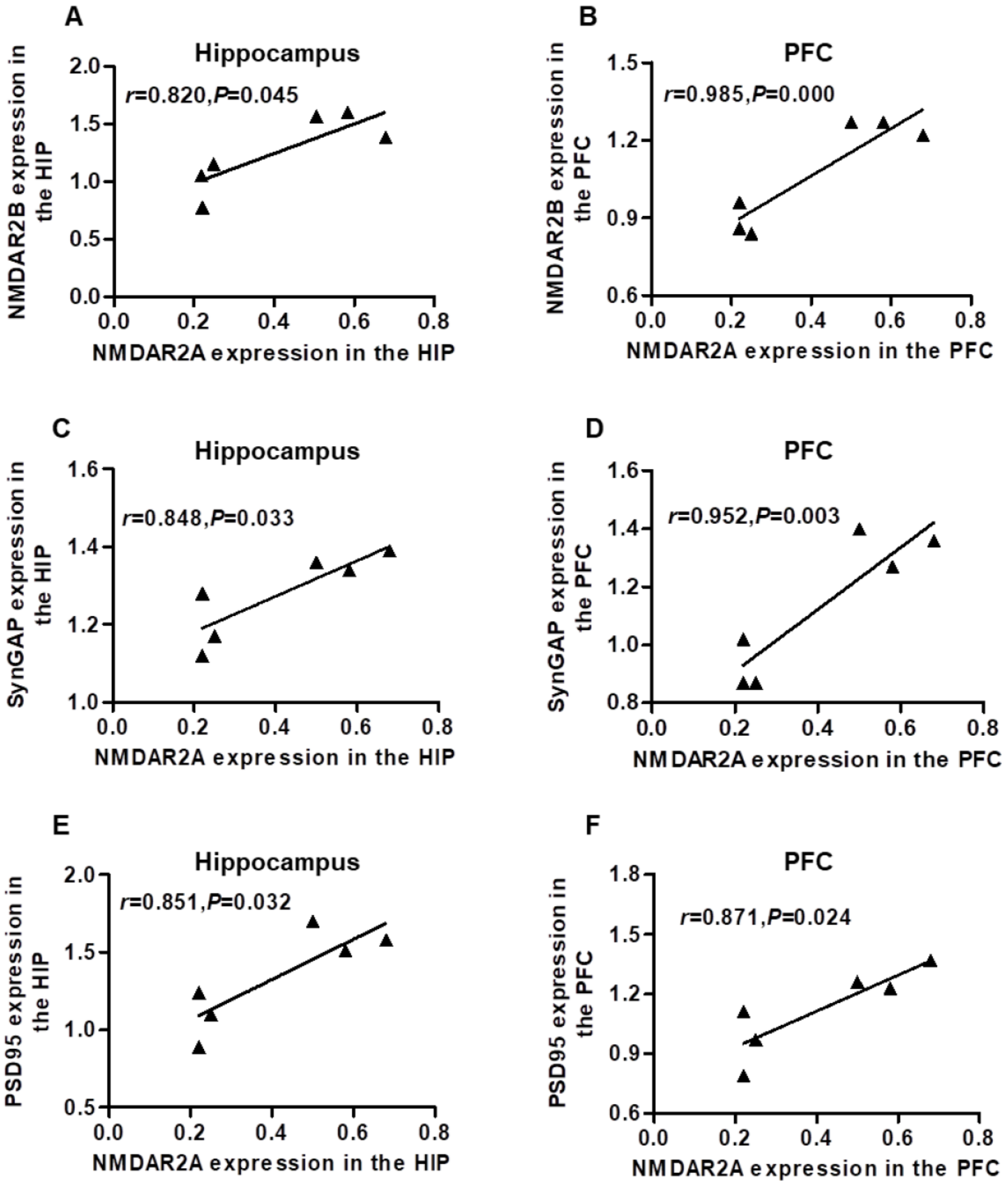


**Supplement 2:** Correlation analysis of the protein expressions between NMDAR2A and NMDAR2B, synGAP, PSD95 in the hippocampus and PFC.

A positive correlation was found between the protein expression of NMDAR2A and NMDAR2B (A, B), synGAP(C, D) and PSD95(E, F) in the hippocampus and PFC.

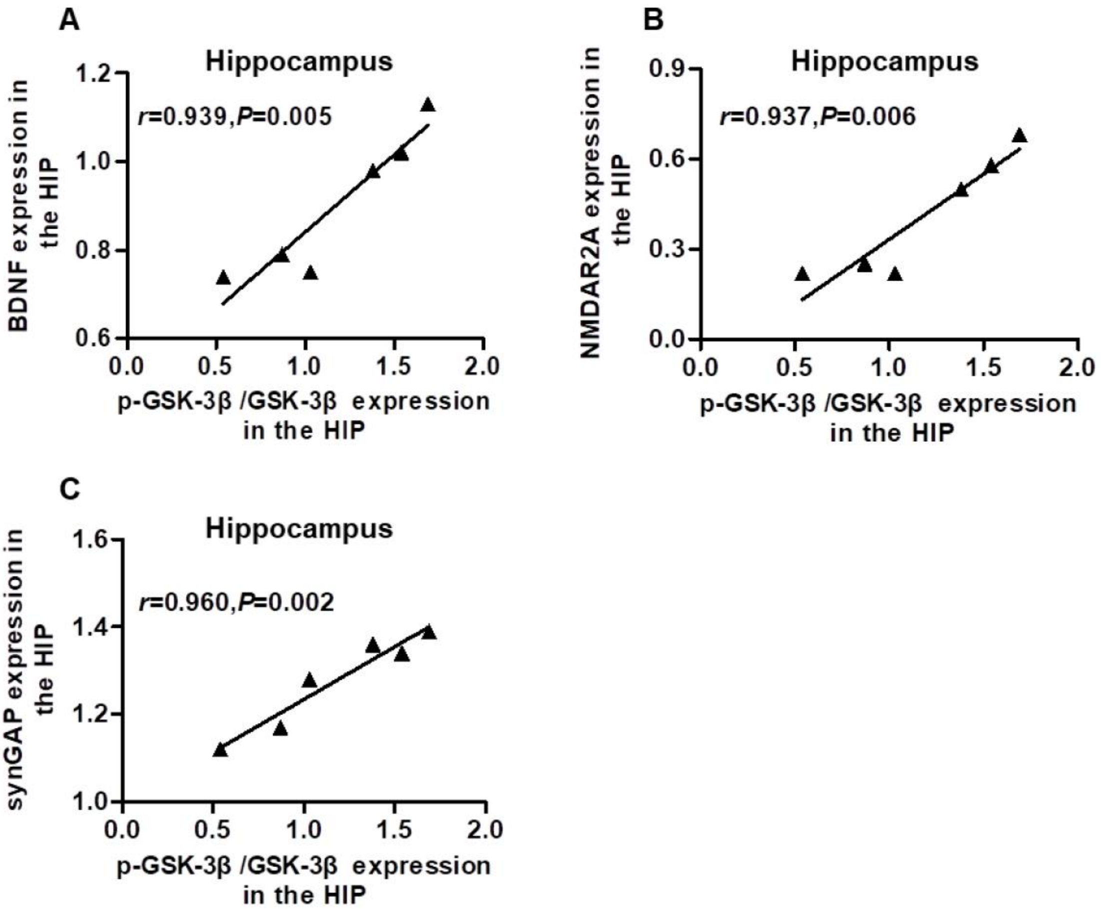


**Supplement 3:** Correlation analysis of the protein expressions between p-GSK-3β/GSK-3β and BDNF, NMDAR2A, synGAP in the hippocampus.

In the hippocampus, p-GSK-3β/GSK-3β expression was positively related to the expression of BDNF (A), NMDAR2A (B), synGAP (C).
